# Skeletal Growth Retardation in Diamond Blackfan Anemia Syndrome Arises From Cell-Autonomous, p53-Dependent Mesenchymal Progenitor Defects Independent of Hematopoietic Failure

**DOI:** 10.64898/2026.11.22.768398

**Authors:** Alexis Bertrand, Jimmy Hom, Yuefeng Tang, Julien Papoin, Theodoros Karnavas, Brian M. Dulmovits, Mushran Khan, Riaz Rao, Benjamin Daniels, Hiren Patel, Emily Hartman, Jedediah Bondy, Morris Edelman, Renaud Touraine, Geneviève Chanoz-Poulard, Gregory Ottenberg, Robert Maynard, Daniel A Grande, Philippe Marambaud, Betsy J Barnes, Anupama Narla, Narla Mohandas, Irene Roberts, Leonard I. Zon, Sébastien Durand, Luanne L. Peters, Lydie Da Costa, Jeffrey M. Lipton, Cheryl L. Ackert-Bicknell, Lionel Blanc

## Abstract

Ribosomopathies are a class of disorders caused by defects in the structure or function of the ribosome and are characterized by tissue-specific abnormalities. Diamond Blackfan anemia syndrome (DBAS) features skeletal anomalies in addition to anemia, but whether these reflect primary mesenchymal dysfunction remains unknown. Here, we demonstrate that skeletal growth retardation in DBAS is cell-autonomous, arising from RPS19 haploinsufficiency in mesenchymal progenitors without leading to hematopoietic failure. RPS19 haploinsufficient mice exhibit shorter limbs due to growth plate disorganization. Single-cell RNA sequencing reveals impaired chondrocyte differentiation, with decreased *Runx2* expression and activation of the p53 pathway across the chondrocyte trajectory. Surprisingly, in vivo translation remains unchanged, arguing against a translational collapse and implying the p53-dependent nucleolar surveillance as main mechanism of growth defect. Accordingly, *Trp53* deletion in RPS19 haploinsufficient mice rescues all skeletal defects in a dose-dependent manner. Importantly, RPL5 haploinsufficiency in mesenchymal progenitors does not result in skeletal defects, establishing ribosomal protein gene specific functions in the mesenchyme. These findings demonstrate that nucleolar surveillance functions as a developmental checkpoint controlling growth plate formation, with direct implications for understanding the tissue-specific manifestations of ribosomopathies.

## INTRODUCTION

Ribosomes are essential organelles involved in protein translation and have long been considered so fundamental to cellular behavior that variations in the biosynthesis or function of these complex macromolecules were considered incompatible with normal biological and cellular function. However, studies have highlighted the ribosome as a cell type-specific regulator of mRNA translation with temporal regulation during development(1–5). The complexity of the eukaryotic ribosome, consisting of four structural ribosomal RNAs and approximately 80 small and large subunit ribosomal proteins (RPs), offers ample opportunity for modifications that permit both temporal and tissue-specific regulation of translation(6, 7). In particular, cell-specific ribosomes have been shown to be variable in their RP composition in specific contexts, across different tissues and at different stages of development(6). This emerging concept of ribosome heterogeneity raises a fundamental question: if different cell types have different ribosomal requirements, do mutations in different RP genes cause different cell type-specific vulnerabilities?

Defects in RP expression or rRNA processing lead to a category of syndromes known as ribosomopathies (8). Diamond Blackfan anemia syndrome (DBAS) is among the best characterized ribosomopathy and is a striking illustration of the tissue-specific consequences of RP haploinsufficiency (9). DBAS is an inherited bone marrow failure syndrome caused by heterozygous mutations in more than 26 RP genes, yet despite the ubiquitous expression and essential function of these genes, disease manifestations are restricted to specific cell types and tissues. Haploinsufficiency of RP genes leads to downstream cellular consequences including impaired translation and activation of the p53 tumor suppressor pathway through nucleolar surveillance(9–11). The prevailing holds that haploinsufficiency of a small subunit RP such as RPS19, the most commonly mutated gene and accounting for approximately 25% of cases, disrupts 40S ribosome assembly. This causes accumulation of free large subunit proteins RPL5 and RPL11 that bind and inhibit the p53 ubiquitin ligase MDM2, leading to p53 stabilization and cell cycle arrest or apoptosis(12).

While the erythroid failure in DBAS has been extensively studied under this framework, DBAS is in fact a multi-system disorder. Beyond red cell aplasia, patients can also present with congenital anomalies that are predominantly skeletal, including craniofacial defects such as oral-facial clefts, vertebral fusions, scapular defects, radial ray anomalies, and shortening of the long bones of the upper limbs(13). Clinical observations suggest that this vulnerability may itself be RP gene-specific: RPS19 mutations are associated with congenital anomalies in approximately 30% of cases while RPL5 mutations are linked to about 70% of cases (14). Finally, patients with DBAS show a predisposition to cancer, and notably solid tumors such as osteosarcoma(15). Despite these distinct and clinically significant skeletal manifestations, the mechanisms leading to skeletal defects and osteosarcoma in DBAS remain largely unknown, as prior studies have predominantly focused on the hematopoietic failure.

A central unresolved question is whether the skeletal and especially growth defects in DBAS represent a primary, cell-autonomous consequence of RP haploinsufficiency in mesenchymal progenitors or whether they arise secondarily to the chronic disease and its treatment. Both the hematopoietic and mesenchymal lineages contribute to bone homeostasis: mesenchymal stromal cells (MSCs) differentiate into osteoblasts and chondrocytes that form bone and cartilage, while hematopoietic stem cells (HSCs) generate the osteoclasts essential for bone resorption(16, 17). DBAS patients experience chronic anemia, prolonged corticosteroid therapy causing steroid-induced bone loss, and transfusion-dependent iron overload — each of these factors may independently impair growth(18). For these reasons, the skeletal defects and low bone mineral density observed in DBAS patients have been attributed to these secondary consequences rather than resulting from a primary autonomous feature of the mesenchymal lineage. Furthermore, whether different RP genes equivalently affect mesenchymal progenitors — or whether the ribosome heterogeneity observed across cell types extends to differential vulnerability of stem cells — has not been addressed.

We previously showed that RP gene identity determines the cellular stage of hematopoietic vulnerability. Indeed, using hematopoietic-specific *Vav-Cre* deletion, we showed that RPS19 haploinsufficiency impairs hematopoietic stem cell function, while RPL5 haploinsufficiency affects erythroid progenitors at a later stage of differentiation(19). Whether this principle of RP gene-specific stem cell vulnerability extends beyond the hematopoietic system to other tissue compartments remains an open question with direct implications for understanding the variable congenital anomalies in DBAS.

Here, using mesenchymal-restricted *Prrx1-Cre* conditional deletion of *Rps19* and *Rpl5*, we demonstrate that skeletal growth retardation in DBAS is a primary, cell-autonomous consequence of RPS19 haploinsufficiency in mesenchymal progenitors, entirely independent of hematopoietic failure. Mechanistically, translation initiation measurements and single-cell RNA sequencing reveal absence of a global translational collapse in favor of a p53-dependent differentiation block. Indeed, MSCs and early chondrocyte progenitors accumulate while mature prehypertrophic and hypertrophic chondrocytes are reduced, indicating that p53-mediated cell cycle arrest prevents mesenchymal progenitors from transiting through the chondrocyte maturation program. This results in dramatic shortening of all limb bones from birth. Importantly, these phenotypes are completely rescued by concomitant p53 deletion in a dose-dependent manner. Strikingly, RPL5 haploinsufficiency in the same mesenchymal progenitors causes no skeletal phenotype, establishing RP gene-specific MSC vulnerability and extending the principle of gene-specific stem cell vulnerability across tissue compartments. We validate these findings in a patient with DBAS carrying a de novo mutation in exon 2 of *RPS19*, who displays impaired rib cartilage. Together, these results reveal that different ribosomal proteins are non-equivalent in their requirements for mesenchymal stem cell function, establishing RP gene-specific stem cell vulnerability as a new paradigm for understanding the tissue-specific manifestations of Diamond-Blackfan anemia syndrome and the broader ribosomopathies.

## RESULTS

### RPS19 haploinsufficiency in the mesenchyme leads to skeletal defects

While numerous studies have reported skeletal defects in patients with a *RPL5* or *RPL11* mutation, these have not been as thoroughly investigated in patients with *RPS19* mutations. A previous case study highlighted growth defects in a stillborn male fetus delivered at ∼37 weeks of gestation with a *de novo* mutation in exon 2 of *RPS19*(20). As demonstrated by x-ray images, this fetus with DBAS presented with shortened long bones of the upper limbs and absence of the deltoid tuberosity of the humerus (**Figure 1A**). To evaluate the cancellous bone architecture of this fetus, we used Masson’s trichrome and Safranin-O staining on the rib bones and compared them to rib bones of a non-hematological stillborn child at the same gestational age (**Figure 1B**). The cancellous bone or bony trabeculae appeared to show reduced connectivity of each trabecular element to adjacent trabeculae in the patient with DBAS. We also noted a relatively thinner calcified matrix centrally within trabeculae and thicker, non-mineralized osteoid matrix on the surface of trabeculae, suggesting a defect in bone mineralization. Finally, we observed reduced cartilage matrix and altered chondrocyte organization. These observations suggested involvement of the mesenchymal lineage in the osteochondral defects of DBAS.

**Figure 1.**
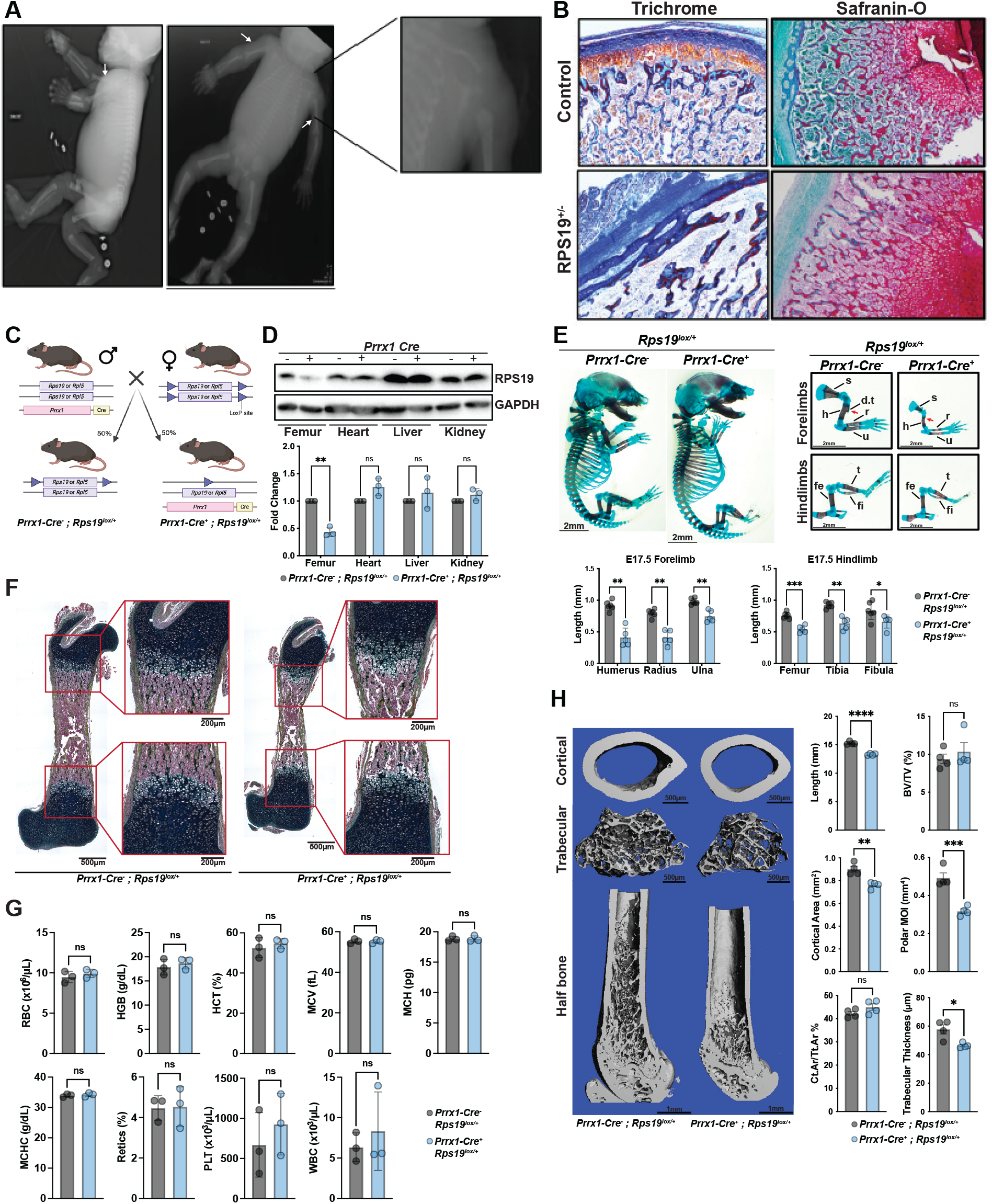
*Prrx1-Cre*-mediated haploinsufficiency of Rps19 recapitulates the human DBAS phenotype. **A.** X-ray images of 37-week-old male fetus heterozygous for a mutation in Exon 2 of *RPS19* who died *in utero.* White arrows denote short proximal upper extremities (inset image) with a posteroanterior view in the top image and a lateral view in the bottom image. **B.** Masson’s Trichrome and Safranin-O staining of rib bones from the DBAS patient and gestational age-matched *RPS19*^+/+^ stillborn fetus. **C.** Schematic representation of the *Prrx1-Cre Rps19* mouse model generated. **D.** Western blot (upper panel) and quantification (bottom panel) of RPS19 from whole cell lysates of different tissues from neonatal *Prrx1-Cre^-^; Rps19^lox/+^* and *Prrx1-Cre^+^*; *Rps19^lox/+^* mice. Fold change relative to *Prrx1-Cre^-^; Rps19^lox/+^* mice and normalized with GAPDH. N=3, ± SD. **E.** Alcian Blue and alizarin red staining of E17.5 *Prrx1-Cre^-^; Rps19^lox/+^*and *Prrx1-Cre^+^; Rps19^lox/+^*mice. Red arrows denote the deltoid tuberosity. Quantification of length of forelimb and hindlimb bones. N=5 ± SD. Scapula (s), humeri (h), deltoid tuberosity (d.t), ulna (u), femur (fe), tibia (t), fibula (fi). **F.** Masson’s Trichrome staining of distal femur from postnatal P2 *Prrx1-Cre^-^; Rps19^lox/+^* and *Prrx1-Cre^+^; Rps19^lox/+^* mice. **G.** Complete blood count of *Prrx1-Cre^-^; Rps19^lox/+^* and *Prrx1-Cre^+^; Rps19^lox/+^* mice at P21. Upper panel: red blood cells (RBC), hemoglobin (HGB), hematocrit (HCT), mean corpuscular volume (MCV), mean corpuscular hemoglobin (MCH). Lower panel: mean corpuscular hemoglobin concentration (MCHC), reticulocytes (Retics), platelets (PLT), white blood cells (WBC) N=3 ± SD. **H.** 3D reconstructed microCT images of cortical bone of the femur at mid-diaphysis, trabecular compartment of the proximal femur, and half-bone from 10-week-old *Prrx1-Cre^-^; Rps19^lox/+^*and *Prrx1-Cre^+^; Rps19^lox/+^*mice and their associated quantification. N=4 ± SD. Bone volume density (BV/TV), moment of inertia (MOI), Cortical area (Ct.Ar), Total area (Tt.Ar). All the *P*-values were calculated using multiple t-tests and adjusted for multiple comparisons using the Holm-Šídàk method. Not significant (ns); *\*P < 0.05; **P < 0.01; ***P < 0.001; ****P < 0.0001*.

To understand if these findings are specific to one clinical case or whether these defects reflect primary mesenchymal dysfunction in DBAS, we generated *Rps19^lox/+^* mice where *Cre* expression is driven by the paired-related homeobox gene-1 (*Prrx1*) promoter, beginning in the mesenchymal stem cell (MSC) and continuing through the osteoblast precursor stage(21) (**Figure 1C**). The *Prrx1-Cre^+^; Rps19^lox/+^* animals were born with a normal Mendelian ratio (33 *Prrx1-Cre^+^; Rps19^lox/+^* live born of 64 total offspring) with a specific deletion of *Rps19* in both endochondral and intramembranous bone as well as in the femoral articular cartilage, as measured at both the genomic (**Figure S1A**) and at the protein level for the long bones (**Figure 1D**). In the femur, the expression levels of RPS19 were lowered by ∼50% (**Figure 1D**) while, as expected, levels in heart, liver and kidney remain identical to those of *Prrx1-Cre^+^; Rps19^+/+^* mice. We did not observe any live *Prrx1-cre; Rps19^lox/lox^* animals, suggesting that *Rps19* is essential for the proper development of the mesenchymal lineage.

Alizarin red and Alcian blue staining of the whole skeleton and the forelimbs at E17.5 and P1 showed hypoplasia of the humerus and the scapula (**Figure 1E, Figure S1B**). These defects were maintained in 10-week-old mice (**Figure S1C**). In addition, the deltoid tuberosity was missing in *Prrx1-Cre^+^; Rps19^lox/+^* (**Figure 1E**, red arrows) in association with significant reduction in humeral and ulnar length (**Figure 1E, bottom panel, Figure S1B**). The hindlimbs of *Prrx1-Cre^+^; Rps19^lox/+^* animals appeared less affected, but we did observe a small, significant decrease in tibial length (**Figure 1E bottom panel, Figure S1B**). Whole-mount skeletal preparations at P2 confirmed generalized limb shortening with altered growth plate architecture (**Figure 1F**). As the *Prrx1-Cre* is known to be active in selected craniofacial mesenchymal lineages, we also examined the calvaria in neonatal mice. Heterozygous mice presented with an increased opening of the anterior fontanelle due to an incomplete growth of the posterior frontal bone; but this defect resolved by P21 (**Figure S1D-E**). Consistent with patients with DBAS harboring a mutation in *RPS19*, we did not observe any incidence of orofacial clefting. Finally, complete blood counts at P21 were normal across all lineages (**Figure 1G**), suggesting that the osteochondral defects occur independently of hematopoietic failure. Together, these data suggest that the contribution of *Rps19* to skeletal development is site-specific, and that our conditional mouse model can serve as model to study the skeletal growth defects observed in the stillborn patient.

We then examined the impact of *Rps19* loss on the young adult cortical and trabecular bone compartments. While 10-week-old mice are still rapidly growing, trabecular bone loss begins at or just before weaning(22). We chose 10 weeks of age as the mice are sexually mature, but not old enough to have experienced substantial maturity-associated cancellous loss in the long bone.

The femur was examined as this is the site typically examined in transgenic mouse models to look for cortical and trabecular bone defects(23). The haploinsufficient femurs of 10-week-old mice were shorter compared to age-matched wild type mice (**Figure 1H**). As expected, this resulted in a smaller cortical bone cross-sectional area at the mid-diaphysis and a highly significant reduction in polar 2nd moment of area (polar moment of inertia, MOI). This latter measure suggests that the femurs of the haploinsufficient mice would be weaker in a twisting breaking configuration. The ratio of the maximum diameter to the minimum diameter of the femur, a measure of roundness, was significantly smaller in the haploinsufficient femurs, likely contributing to the lower polar moment. As such, the bones of the haploinsufficient mice were not only anatomically smaller overall but also demonstrated a different overall morphology beyond the gross anatomical differences noted above (**Figure 1E, Figure S1B**). While we observed no difference in the trabecular bone volume fraction (the ratio of trabecular bone volume to the total volume of the cancellous compartment), there was a significant difference in trabecular thickness, which is reminiscent of the trabecular configuration we observed for the stillborn fetus. Because *Prrx1*-cre is known not to be expressed in the axial skeleton(24), we could not determine whether *Rps19* haploinsufficiency impacts the axial skeleton in our model.

Altogether, these results suggest that the skeletal defects observed in Rps19 haploinsufficiency are due to mesenchymal failure leading to the disorganization of the growth plates, without leading to hematopoietic failure.

### RPS19 deficient mice present a chondrocyte lineage progression defect

To test this hypothesis and further define the molecular mechanisms underlying the skeletal growth retardation due to Rps19 haploinsufficiency, we performed single-cell RNA sequencing (scRNAseq) on E17.5 forelimbs and hindlimbs from two *Prrx1-Cre; Rps19^lox/+^* and two control littermates (**Figure 2A**). After quality filtering and balanced down-sampling (**Figure S2A-B**), the results were analyzed using an unsupervised approach as previously described(25). Cluster identities were assigned based on the expression of the most highly expressed cluster-specific marker genes, enabling us to distinguish 24 different populations (**Figure 2B**). All expected mesenchymal populations were observed, including MSCs, chondrocyte progenitors, and the full chondrocyte differentiating stages from resting through proliferative, prehypertrophic, and hypertrophic stages, as well as early skeletal and cells of hematopoietic origin (**Figure 2B**). Comparisons of the clusters between *Rps19^lox/+^* and control littermates based on gene expression profiles revealed that most alterations were observed in the MSC and the chondrocyte populations frequencies (**Figure S2C, Figure S3A**). This suggests that the MSC and chondrocyte lineage are more affected during development of the limbs due to the loss of one allele of *Rps19*. To test this hypothesis further, we performed an analysis of the chondrocyte trajectory which revealed a striking shift in composition in *Rps19^lox/+^* limbs compared to the littermate controls (**Figure 2C-E**). Indeed, in both forelimbs and hindlimbs we observed a relative accumulation of the MSC population, while the more engaged chondrocytic populations were decreased (**Figure 2D, Figure S3B**). The phenotype was stronger in the forelimbs, consistent with their one-day earlier developmental timing(21, 26). This is consistent with the clinical presentation in the patient with DBAS and strongly suggest that the phenotype is a true biological effect rather than a *cre* artifact. Slingshot analyses confirmed that Rps19 haploinsufficient cells showed impaired progression through the chondrocyte maturation sequence in both forelimbs and hindlimbs (**Figure 2E, Figure S3C**). Cluster-specific gene expression analysis provided molecular insights into the impairment in differentiation (**Figure 2F**). In forelimb MSCs, the master skeletal differentiation transcription factor *Runx2* was significantly reduced, while the chondrogenic identity factor *Sox9* was unchanged, indicating that MSCs preserve their lineage identity but partially lose the capacity to initiate the differentiation program. Consistent with impaired commitment, the chondrocyte matrix gene *Col2a1* was reduced in MSCs and chondrocyte progenitors, and *Sox9* expression was decreased in chondrocyte progenitors. In contrast, *Runx2* expression was elevated in chondrocyte progenitors and hypertrophic chondrocytes, suggesting that the few cells that do transit into the development program exhibit compensatory upregulation. The hypertrophic marker *Col10a1* was reduced in forelimb hypertrophic chondrocytes, consistent with depletion of the terminal chondrocyte population, and a similar trend was observed in the hindlimbs.

**Figure 2.**
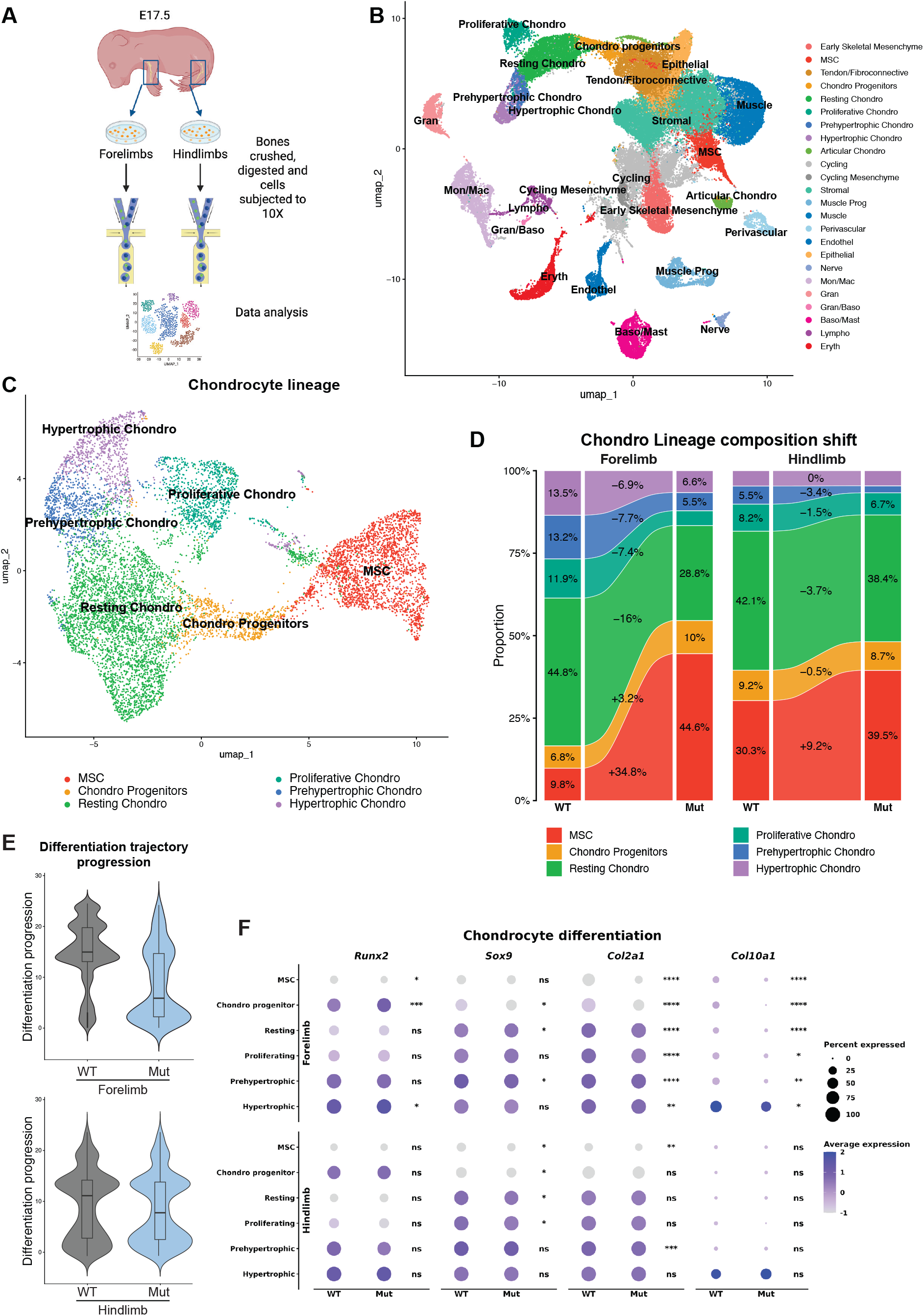
scRNAseq analyses unravel a chondro-lineage progression defect in the mesenchymal Rps19 haploinsufficient model. **A.** Schematic representation of the single cell RNA sequencing strategy. **B.** scRNA-seq UMAP of integrated limbs from control (*Prrx1^-^*) and mutant (*Prrx1^+^*) *Rps19^lox/+^* cells with clusters identified by marker genes. **C.** Refined UMAP focused on the chondro-lineage determined by slingshot. **D.** Alleviate-plot representation of the chondro-lineage proportion in forelimb and hindlimb from control (*Prrx1^-^*) and mutant (*Prrx1^+^*) *Rps19^lox/+^* cells. **E.** Differentiation progression across the chondro-lineage in forelimb and hindlimb from control (*Prrx1^-^*) and mutant (*Prrx1^+^*) *Rps19^lox/+^*cells. **F.** Dot plots showing the expression of *Runx2*, *Sox9*, *Col2a1*, and *Col10a1* across chondrocyte populations in forelimbs (top) and hindlimbs (bottom). Dot size represents the percentage of cells expressing each gene, and color indicates scaled average expression. WT and mutant cells are compared within each cell population. *P*-values were calculated using a Wilcoxon rank-sum test and adjusted for multiple comparisons using the Benjamini-Hochberg method. *ns, not significant; *P < 0.05; **P < 0.01; ***P < 0.001; ****P < 0.0001*.

Together, these data suggest that this failure of MSCs to generate adequate numbers of differentiated chondrocytes could lead to the impaired growth plate architecture and longitudinal bone growth observed from E17.5.

### Cell-autonomous proliferative and differentiation delays in Rps19 haploinsufficient limbs are a consequence of p53 activation in the chondrocyte lineage

To complement the proportional compositional analysis from scRNAseq with quantitation of absolute cell numbers, we performed flow cytometry on E17.5 limb mesenchyme. While the total cell number was similar between control versus *Rps19^lox/+^* limbs, MSCs (immunophenotypically defined as CD31⁻Lin⁻Sca1⁺CD51⁺ (25)) were significantly reduced in absolute number in hindlimbs (*p=0.0002*) with a trend noted in forelimbs (*p=0.13*) (**Figure 3A-B**). This absolute reduction, occurring alongside the proportional enrichment of MSCs within the chondrocyte trajectory observed by scRNAseq (**Figure 2E**), indicates that while the late-stage chondrocyte populations are depleted more severely than MSCs — causing their relative enrichment — the overall mesenchymal progenitor pool is also contracting. We recently demonstrated that the erythroid failure in Rps19 haploinsufficiency begins at the HSPC stages(19) and it has been previously shown that there is crosstalk between the hematopoietic and mesenchymal compartments(27). Thus, we quantified HSPCs in the bone marrow of Rps19 haploinsufficient and control littermates at E17.5 (**Figure 3C**). Consistent with the complete blood counts in the adult mice (**Figure 1H**), the numbers of HSPCs were similar between the two groups in both the forelimbs and the hindlimbs (**Figure 3D**), strengthening the notion that the growth defects are cell autonomous.

**Figure 3.**
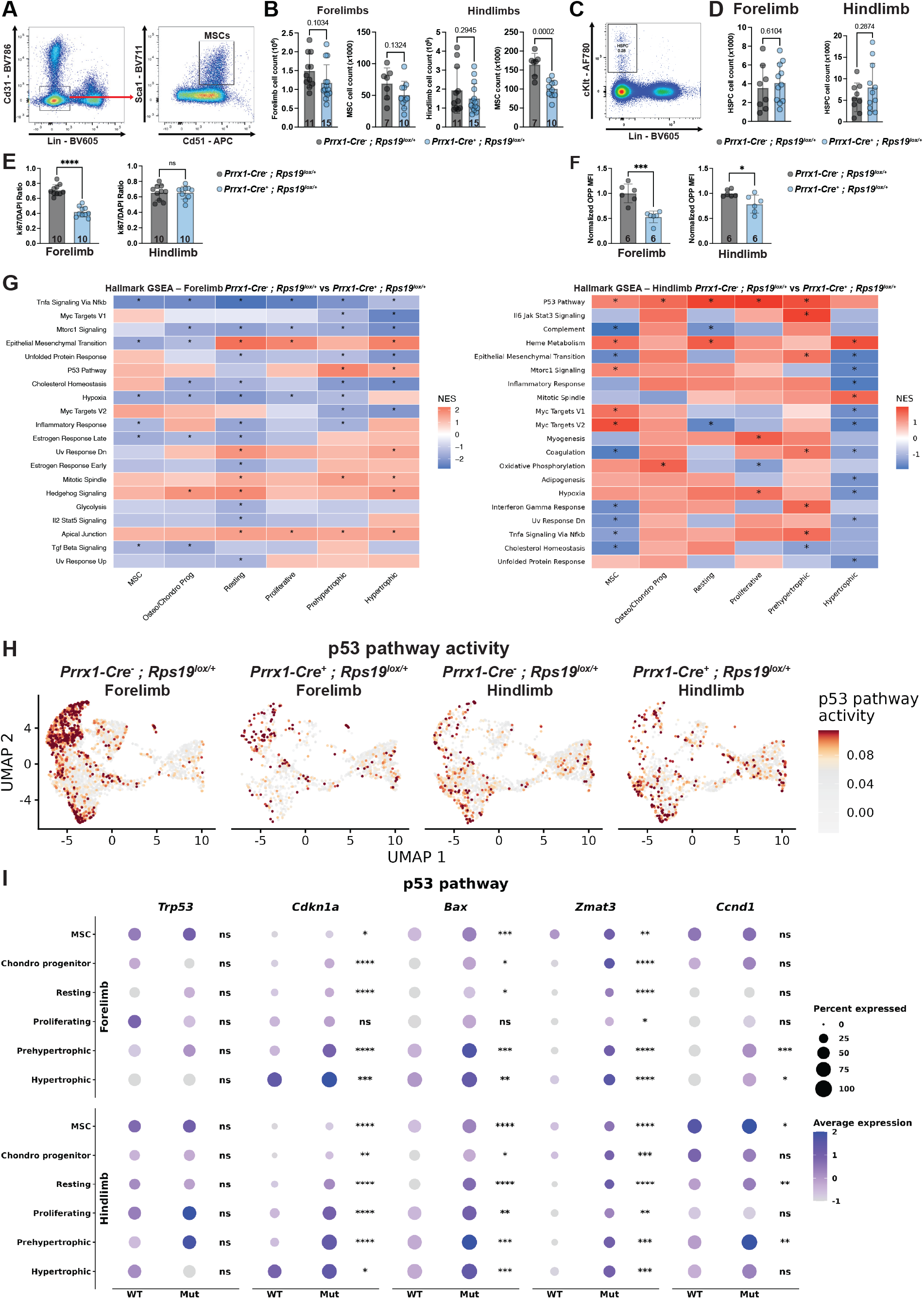
Rps19 haploinsufficiency disrupts mesenchymal progenitor homeostasis and activates p53 signaling during skeletal development. **A.** Representative flow cytometry gating strategy for the identification of mesenchymal stromal cells (MSCs), defined as Lin⁻CD31⁻CD51⁺Sca1⁺ cells, in embryonic limbs. B. Total limb cell and MSC count in forelimb (left) and hindlimb (right) in control (*Prrx1^-^*) and mutant (*Prrx1^+^*) *Rps19^lox/+^* E17.5 embryonic limb. **C.** Representative flow cytometry gating strategy for the identification of hematopoietic stem and progenitors cells (HSPCs) defined as Lin⁻cKit^+^ cells, in E17.5 bone marrow. **D.** Quantification of HSPCs number in forelimb and hindlimb in control (*Prrx1^-^*) and mutant (*Prrx1^+^*) *Rps19^lox/+^* E17.5 bone marrow. **E.** *In vitro* MSC proliferation assessment by Ki67 expression. **F.** *In vitro* MSC protein synthesis measurement by O-propargyl-puromycin (OPP) incorporation. OPP mean fluorescence intensity (MFI) was normalized to the control group. **G.** Hallmark Gene Set Enrichment Analysis (GSEA) comparing WT (*Prrx1^-^*) and mutant (*Prrx1^+^*) *Rps19^lox/+^* cells across MSC and chondrocyte populations in forelimbs (left) and hindlimbs (right). Heatmaps show normalized enrichment scores (NES) for selected Hallmark pathways. Asterisks indicate pathways with significant GSEA results. **H.** UMAP visualization of p53 pathway activity along the chondro-lineage in control (*Prrx1^-^*) and mutant (*Prrx1^+^*) *Rps19^lox/+^* forelimbs and hindlimbs cells. Cells are colored according to their p53 pathway activity score. **I.** Dot plots showing the expression of *Trp53*, *Cdkn1a*, *Bax*, *Zmat3* and *Ccnd1* across chondrocyte populations in forelimbs (top) and hindlimbs (bottom). Dot size represents the percentage of cells expressing each gene, and color indicates scaled average expression. WT and mutant cells are compared within each cell population. *P*-values were calculated using a Wilcoxon rank-sum test and adjusted for multiple comparisons using the Benjamini-Hochberg method. *ns, not significant; *P < 0.05; **P < 0.01; ***P < 0.001; ****P < 0.0001.* The number of replicates is indicated under each graph.

The decrease in the number of immunophenotypically-defined MSCs in Rps19 haploinsufficient mice led us to interrogate their proliferative and overall translation initiation capacity. MSCs were isolated from the forelimbs and hindlimbs of 3-4-week-old mice and cultured in vitro. Using Ki67 as proliferation marker, we observed a specific decrease in the proliferation of MSCs isolated from the forelimbs, while it remained unaffected in hindlimbs-derived MSCs (**Figure 3E**). To measure global translation, we incubated MSCs with O-propargyl-Puromycin (OPP) and then measured the fluorescence intensity by flow cytometry(28). We observed a global decrease in protein synthesis in the *Rps19^lox/+^* MSCs independent of their bone origin (**Figure 3F**). These data suggest that MSCs derived from Rps19 haploinsufficient forelimbs have decreased proliferation and translation capacity. However, when the same measurements were performed *ex vivo* using cells directly isolated from the limbs of haploinsufficient mice, we did not observe changes in the cell cycle (**Figure S4A**) or a decrease in OPP incorporation (**Figure S4B**) suggesting that the *in vivo* defect is buffered by the microenvironment.

Numerous studies have demonstrated impaired proliferation in Rps19 haploinsufficiency and ribosomopathies in general, linking them to the p53 pathway in the hematopoietic compartment(12, 19, 29, 30). However, much less is known regarding the role of p53 in the mechanism of abnormal growth plate architecture and skeletal defects in DBAS. Gene set enrichment analysis (GSEA) across the chondrocyte lineage revealed increased and sustained activation of the p53 pathway in Rps19 haploinsufficient cells compared to littermate controls in both forelimbs and hindlimbs (**Figure 3G**). Since Trp53 activation regulates proliferation and hypertrophy during growth plate maturation, we next assessed p53 pathway activity score across the chondrocyte lineage in *Rps19^lox/+^* and littermate controls (**Figure 3H**). While wild type cells showed the expected increase in p53 activity during terminal chondrogenesis, mutant limbs displayed aberrant p53 activity in early progenitors. At the individual genes level, *Cdkn1a* and *Zmat3* were elevated predominantly in prehypertrophic and hypertrophic chondrocytes (**Figure 3I, Figure S3**), consistent with p53-mediated cell cycle arrest at late stages of chondrocyte maturation. While *Bax* showed modest elevation in some populations, the predominant cellular response appeared to be cell cycle arrest rather than apoptosis, consistent with unchanged *Bax* expression in MSCs. Together, these results suggest that activation of the p53 pathway in the chondrocytic lineage drives delayed cell-autonomous proliferative and differentiation in Rps19 haploinsufficient limbs.

### Growth plate defects in RPS19 haploinsufficiency are p53-mediated

To directly assess the contribution of *Trp53* to the growth defects observed in the Rps19 haploinsufficient model, we bred *Prrx1-Cre^+^; Rps19^lox/+^* to *Trp53^fl/fl^* mice. There was a dose-dependent rescue of the whole-body size at birth (**Figure 4A-B**) in association with *Trp53* dose-dependent restoration of the humeral and scapular morphology, and a complete correction of the cranial defects observed at birth (**Figure S5**). Concomitantly, p53 deletion restored growth plate organization in a dose-dependent manner, with heterozygous p53 deletion providing partial restoration and homozygous p53 deletion returning the growth plate to near-normal architecture (**Figure 4C**). This rescue was further confirmed at the cellular level, with changes in COL2A1 and COL10A1 immunofluorescence staining in the femurs (**Figure 4D-E**). Indeed, the proliferative zone marked by COL2A1 expression and the hypertrophic zone marked by COL10A1 expression were both altered in mutant growth plates compared to controls, and the pattern progressively returned to its physiological configuration following removal of one or two alleles of *trp53*. Interestingly, further analyses of the growth plates at P21 showed that the alterations in COL2A1 and COL10A1 were less pronounced than at P1 in the Rps19 haploinsufficient femurs (**Figure 4F-G**), suggesting that the defects observed during embryogenesis and at birth are actually reflective of a developmental delay rather than a true defect in the mutant mice.

**Figure 4.**
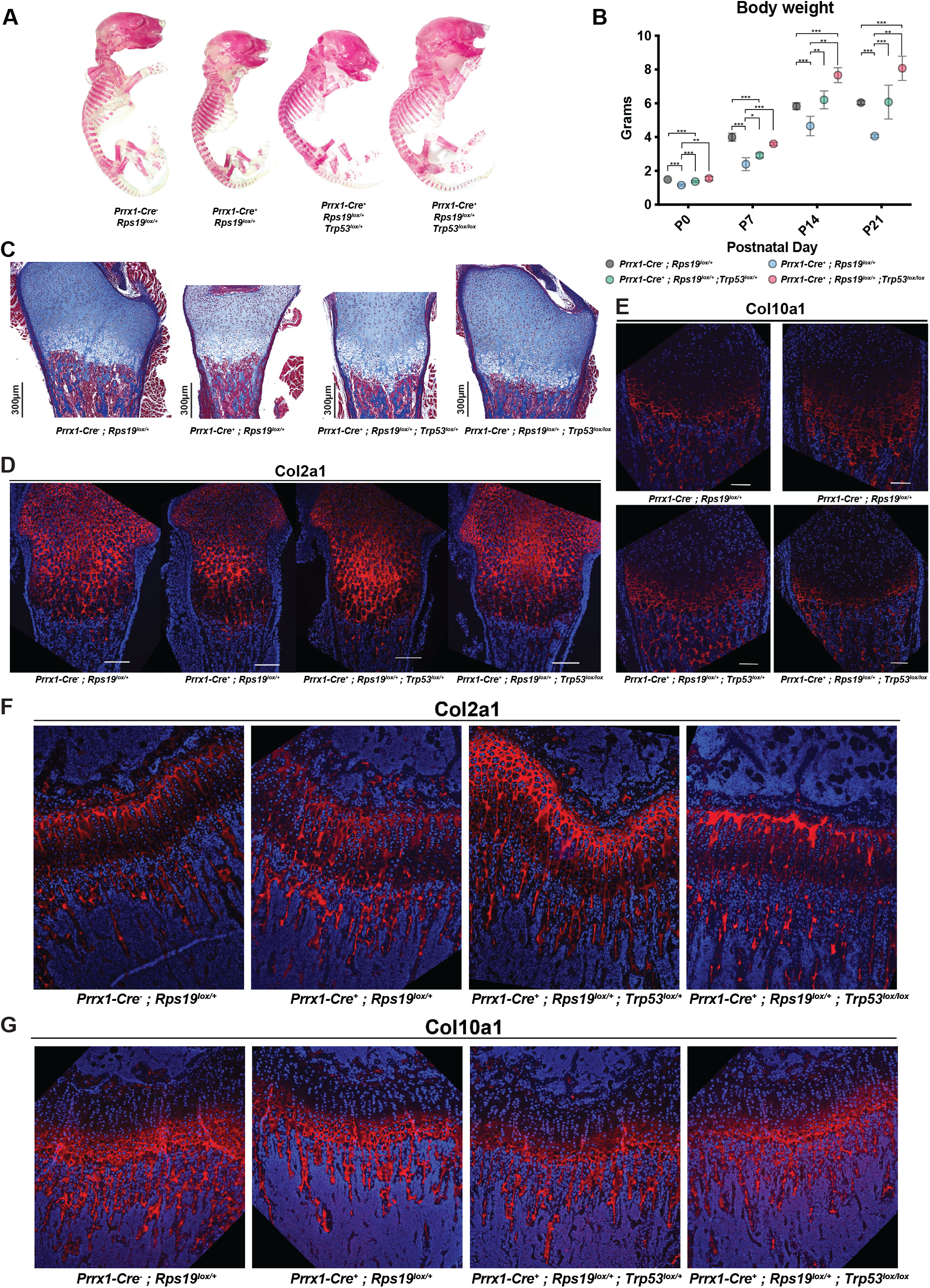
*Trp53* depletion rescues growth plate defects induced by Rps19 haploinsufficiency in a dose-dependent manner. **A.** Alizarin red whole mount skeletal staining of neonatal pups (P1). **B.** Body weight measurement measured weekly from P0 to P21. N=6. (ANOVA *\*P<0.05, **P<0.01, ***P<0.001*). **C.** Masson’s trichrome staining of distal femur from newborn P1 pups. **D.** Representative COL2A1 immunofluorescence staining of distal femur from newborn P1 pups. DAPI=blue; COL2A1=Red. **E.** Representative COL10A1 immunofluorescence staining of distal femur from newborn P1 pups. DAPI=blue; COL10A1=Red. **F.** Representative COL2A1 immunofluorescence staining of distal femur from three weeks mice. DAPI=blue; COL2A1=Red. **G.** Representative COL10A1 immunofluorescence staining of distal femur from three weeks mice. DAPI=blue; COL10A1=Red.

### Mice with RPL5 haploinsufficiency in mesenchymal stem and progenitor cells present normal skeletal growth

To determine whether the skeletal phenotype is a general consequence of ribosomal protein haploinsufficiency in mesenchymal progenitors or reflects gene-specific vulnerability, we generated *Prrx1-Cre; Rpl5^lox/+^* mice using a validated *Rpl5* conditional allele that leads to a DBAS phenotype in the hematopoietic compartment(19). Cre-mediated excision of the *Rpl5* floxed allele was confirmed by western blot in limb tissues (**Figure 5A**). Strikingly, *Prrx1-Cre^+^; Rpl5^lox/+^* mice displayed no apparent altered skeletal phenotype. Body weight, complete blood counts, and gross skeletal morphology were indistinguishable from wild-type littermates at all time points examined (**Figure 5B-D**). Finally, we measured the number of immunophenotypically defined MSC at E17.5 as we did in the case of Rps19 haploinsufficient mice **(Figure 5E)**. The total number of cells in the limbs was similar between *Prrx1-Cre^+^; Rpl5^lox/+^*and control littermates, and MSC numbers showed no difference between the two groups, unlike what was observed in the case of Rps19-haploinsufficient mice. Thus, deletion of one allele of Rpl5 in the MSCs did not lead to growth retardation.

**Figure 5.**
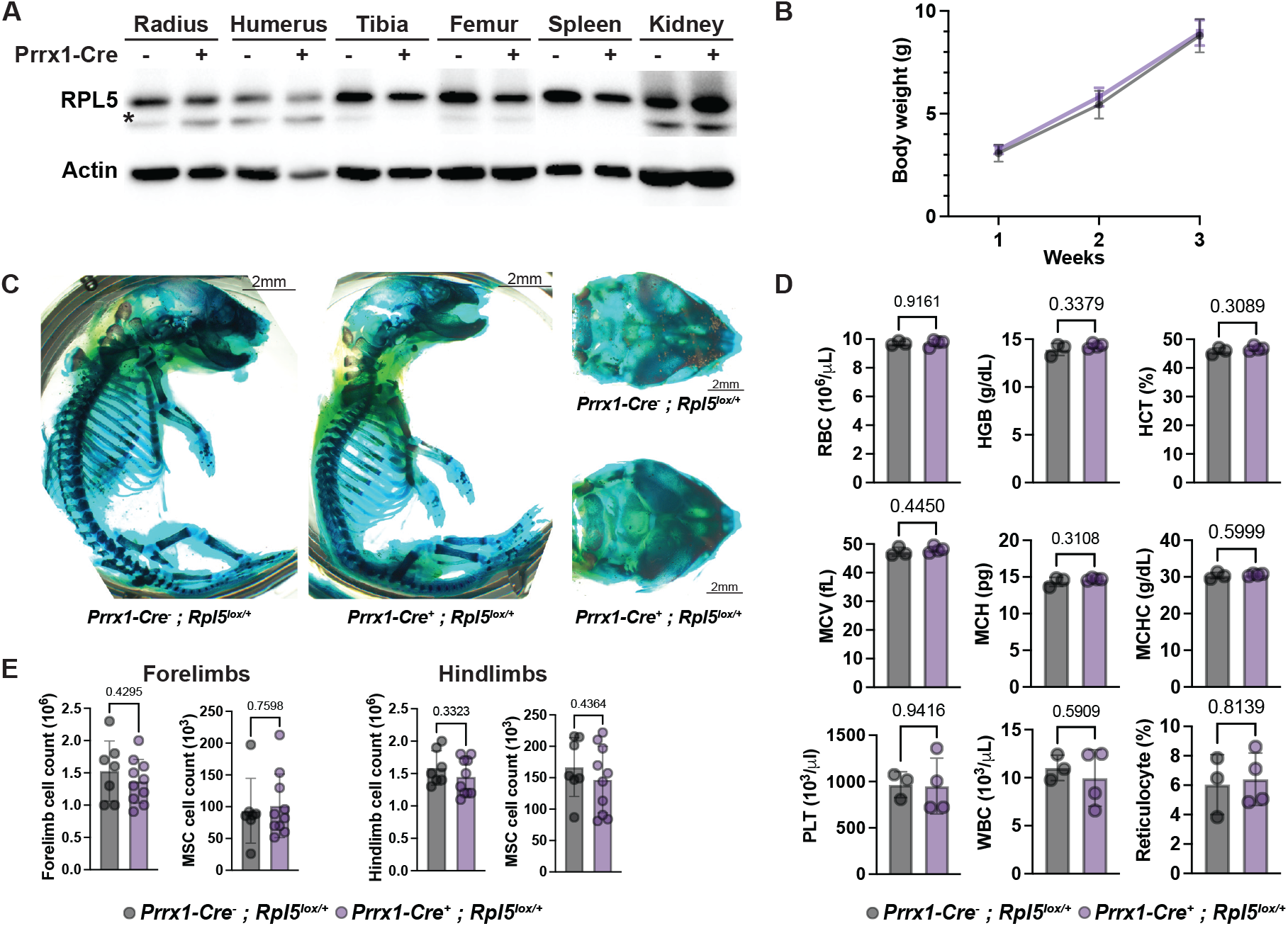
*Prrx1-Cre^+^; Rpl5^lox/+^* mice do not present growth defects. **A.** Western blot of RPL5 and GAPDH from whole cell lysates of different tissues from *Prrx1-Cre^-^; Rpl5^lox/+^* and *Prrx1-Cre^+^*; *Rpl5^lox/+^* mice. Asterisk denotes nonspecific band **B.** Body weight measurement measured weekly from P0 to P21. N=5-9 ± SD. (2-way ANOVA, *ns, not significant*). **C.** Alcian Blue staining of P1 *Prrx1-Cre^-^; Rpl5^lox/+^*and *Prrx1-Cre^+^; Rpl5^lox/+^*mice. **D.** Complete blood count of 6-weeks old *Prrx1-Cre^-^; Rpl5^lox/+^* and *Prrx1-Cre^+^; Rpl5^lox/+^* mice. Upper panel: red blood cells (RBC), hemoglobin (HGB), hematocrit (HCT). Middle panel: mean corpuscular volume (MCV), mean corpuscular hemoglobin (MCH), mean corpuscular hemoglobin concentration (MCHC). Lower panel: platelets (PLT), white blood cells (WBC), reticulocytes (Retics). N=3-4 ± SD. **E**. Total limb cell and MSC count in forelimb (left) and hindlimb (right) in control (*Prrx1^-^*) and mutant (*Prrx1^+^*) *Rpl5^lox/+^* E17.5 embryonic limb. Except for (**B**) all the *P*-values were calculated using multiple t-tests and adjusted for multiple comparisons using the Holm-Šídàk method.

Together, these results suggest that ribosomal protein haploinsufficiency differentially impacts MSCs, as it does in the hematopoietic compartment(19).

## DISCUSSION

In this study, we demonstrate that skeletal growth retardation in a murine model of DBAS is a primary, cell-autonomous consequence of *RPS19* haploinsufficiency in mesenchymal progenitors, entirely independent of hematopoietic failure. Single-cell RNA sequencing reveals a differentiation delay at the MSC-to-chondrocyte transition, with MSCs accumulating while all downstream chondrocyte populations are depleted. This differentiation delay is driven by p53 activation, rescued by its deletion in a dose-dependent manner, and appears specific to RPS19. Although growth plate architecture partially normalizes by P21, the persistent bone shortening at 10 weeks indicates that the developmental delay during the critical embryonic and early postnatal growth period produces an irreversible growth deficit. These findings establish ribosomal protein gene-specific mesenchymal stem cell vulnerability as a determinant of skeletal growth defects in DBAS.

At the molecular level, RPS19 haploinsufficiency impairs chondrocyte differentiation at two levels along the differentiation trajectory. In MSCs, *Runx2* expression was reduced while *Sox9*’s was maintained, indicating that MSCs preserve their lineage identity but lose differentiation capacity(31, 32). This finding is consistent with established evidence that p53 directly represses *Runx2* transcription(33), providing a direct molecular link between nucleolar surveillance and the differentiation block. Downstream, the p53 target *Cdkn1a/p21* was elevated in prehypertrophic and hypertrophic chondrocytes, arresting cells that had already entered the maturation program. The convergence of impaired differentiation initiation and delayed maturation explains the scRNAseq composition shift – MSC relative accumulation with depletion of all downstream populations – and the growth plate disorganization observed histologically. Notably, global protein translation was unchanged *in vivo* despite being reduced in cultured MSCs, indicating that the skeletal phenotype reflects p53-dependent developmental surveillance rather than translational insufficiency, and that niche-derived signals may partially buffer translational defects.

The more severe forelimb growth phenotype — with greater scRNAseq composition shifts and humerus fragility precluding microCT analysis at 10 weeks — is consistent with the clinical presentation of DBAS, in which upper extremity anomalies are more prevalent than lower limb defects(34). While the earlier onset of *Prrx1* expression in forelimbs (E9.5 vs E10.5 for hindlimbs) could contribute through longer exposure to Rps19 haploinsufficiency, the clinical concordance with germline DBAS mutations suggests intrinsic differences in vulnerability between forelimb and hindlimb mesenchyme. This anatomic site-specific vulnerability may reflect differences in proliferative demand, niche signaling, or ribosome composition between forelimb and hindlimb mesenchymal progenitors and warrants further investigation.

Our study further establishes the cell-autonomous nature of skeletal growth defects in DBAS. The *Prrx1-Cre^+^; Rps19^lox/+^*mice develop severe skeletal growth retardation with normal hematopoiesis. Reciprocally, hematopoietic-specific *Vav-iCre^+^; Rps19^lox/+^*mice develop anemia and hematopoietic disorders without primary skeletal defects, except for overall growth retardation secondary to the anemia(19). This reciprocal independence has direct clinical implications. Indeed, skeletal growth abnormalities in DBAS will not be corrected by gene therapy or hematopoietic stem cell transplantation and require independent monitoring and intervention. Furthermore, the skeletal microenvironment is essential for hematopoietic stem cell maintenance(16, 17, 27), yet the *Prrx1-Cre* model shows that mesenchymal Rps19 haploinsufficiency does not impair the bone marrow niche sufficiently to cause hematopoietic failure, suggesting that niche function is maintained despite the skeletal growth delay.

The absence of any skeletal phenotype in *Prrx1-Cre^+^; Rpl5^lox/+^* mice is striking. Combined with our previous demonstration that RPS19 and RPL5 haploinsufficiency affect distinct stages of hematopoiesis(19) a coherent framework emerges, in which RPS19 haploinsufficiency preferentially affects early stem and progenitor cells across tissue compartments, while RPL5 haploinsufficiency affects more committed progenitors within specific lineages. Congenital anomalies are rarely reported in patients with *RPS19* mutations, with skeletal involvement resulting from impaired growth while historically, congenital anomalies are attributed primarily to RPL5 and RPL11 genotypes(14) likely resulting from overactivation of growth signals. However, our finding that RPS19 haploinsufficiency profoundly impairs the mesenchymal stem cell compartment during development, consistent with its effects on hematopoietic stem cells(19, 35), suggests that the prevalence of skeletal involvement resulting from growth impairment in RPS19-mutant DBA is likely underrecognized. Furthermore, the severity of the skeletal phenotype in our model, combined with the impaired rib cartilage observed in a stillborn patient carrying an *RPS19* exon 2 deletion raise the possibility that the most severely growth retarded *RPS19*-mutant pregnancies result in fetal loss before clinical diagnosis, creating an ascertainment bias that underestimates the full spectrum of skeletal involvement.

The accumulation of p53-activated, differentiation-arrested MSCs in Rps19-haploinsufficient growth plates may also bear on the predisposition to osteosarcoma observed in DBAS patients(15). Chronically arrested progenitors that are viable but unable to differentiate represent a potential reservoir for malignant transformation; a secondary loss of p53 function, through somatic mutation or epigenetic silencing, would simultaneously release the differentiation block and remove tumor suppression, potentially giving rise to mesenchymal malignancies. The dose-dependent rescue we observe with p53 deletion illustrates the inherent tension between developmental rescue and tumor surveillance in therapeutic targeting of the p53 axis.

More broadly, this rescue by p53 suggests a release of the nucleolar surveillance, further indicating that nucleolar surveillance functions as a developmental checkpoint. In this context, restricting MSC differentiation through *Runx2* repression without requiring translational collapse suggests that the p53-mediated quality control response, rather than translational insufficiency per se, may be the common pathogenic mechanism underlying tissue-specific defects in ribosomopathies. The identity of the ribosomal protein affected would then determine which surveillance response is activated in which cell type, producing the characteristic genotype-phenotype correlations observed clinically.

In conclusion, skeletal growth retardation in DBA arises from a p53-dependent differentiation delay in mesenchymal progenitors, in which *Runx2* repression decreases MSC differentiation initiation and p21-mediated arrest depletes late-stage chondrocytes, collapsing the growth plate. This mechanism is cell-autonomous, independent of hematopoietic failure, and specific to RPS19. Indeed, RPL5 haploinsufficiency does not affect the same cell population resulting in mechanistically different characteristic skeletal anomalies. The identification of p53-mediated nucleolar surveillance as a developmental checkpoint governing mesenchymal stem cell fate opens a new avenue for understanding how mutations in ubiquitously expressed ribosomal protein genes cause tissue-specific developmental disorders and establishes the foundation for gene-specific therapeutic and monitoring strategies in DBAS.

## METHODS

### Sex as Biological Variable

Human studies: The patient with DBAS was a male fetus. The control was deidentified and the sex was unknown.

Murine studies: *Prrx-Cre^+^* male mice were used for breeding to avoid the known potential for germline allele excision when *Prrx1*-*Cre* is passed from the dam(21). Embryos and pups from both sexes were analyzed, no differences were observed.

### Antibodies

| CD4--BV605 | **Biolegend** | **Cat# 100451** |
|---|---|---|
| CD8a-BV6-5 | Biolegend | Cat# 100744 |
| Ter119-BV605 | Biolegend | Cat# 116239 |
| CD45R/B220-BV605 | Biolegend | Cat# 103224 |
| Ly-6G/Ly-6C-BV605 | Biolegend | Cat# 108440 |
| C-Kit-APC-eFluor780 | eBioscience | Cat# 47-1171-82 |
| Sca-1-BV711 | Thermo Science | Cat# 11-5981-82 |
| CD51-PE | Thermo Science | Cat# 12-0512-81 |
| CD51-APC | Elabscience | Cat# E-AB-F1235UE |
| CD45-BV605 | Biolegend | Cat# 103155 |
| CD31-BV785 | Biolegend | Cat# 102435 |
| Ki67-PE | Biolegend | Cat# 652404 |
| GAPDH-HRP | CST | Cat# 8884 |
| P53 | CST | Cat# 2524 |
| Rps19 | Bethyl Labs | Cat# A304-002A |
| Rpl5 | Abcam | Cat# ab86863 |
| Actin-B | CST | Cat# 4970 |
| Col1A1 | CST | Cat# 84336 |
| MMP9 | Abcam | Cat# ab38898 |
| **Chemicals, peptides, and recombinant proteins** |  |  |
| Fetal bovine serum | **Cytiva** | **Cat# SH30071.03** |
| IMDM | Fisher Scientific | Cat# 12-440-079 |
| BSA | Fisher Scientific | Cat# BP9703100 |
| PBS | Cytiva | Cat# SH30256.LS |
| RIPA buffer | Thermo Fisher Scientific | Cat# 89900 |
| Precision Plus Protein™ Dual Color Standards | BioRad | Cat# 1610374 |
| 100 bp DNA Ladder | NEB | Cat# N3231L |
| Proteinase K | Lamda Biotech | Cat# DB0451-10 |
| O-Propargyl-Puromycin | MedChem Express | Cat# HY-15680 |
| Phosphatase Inhibitor Cocktail | Bio Basic | Cat# PL017 |
| Protease Inhibitor Cocktail | Bimake | Cat# B14002 |
| DAPI | Sigma-Aldrich | Cat# D9542-10mg |
| Poloxamer 188 | Sigma-Aldrich | Cat# P5556-100ML |
| Medium 199 | Fisher Scientific | Cat# SH30253.01 |
| Collagenase Type 2 | Worthington Biochemical | Cat# LS004177 |
| Alcian Blue | Thermo Scientific | Cat# J60122-09 |
| Alizarin Red S | Sigma-Aldrich | Cat# A5533-10mg |
| Glycerol | Sigma-Aldrich | Cat# G7893-4L |

### Critical commercial assays

Chromium Next GEM Single Cell 3’ Kit (10X Genomics, #1000268); Chromium Next GEM Chip G Single Cell Kit (10X Genomics, #1000120); Dual Index Kit TT, Set A (10X Genomics, #1000215).

### Deposited data

Mouse single cell RNAseq (This paper).

### Transgenic mice

*Prrx1*-*Cre* (JAX #005584) and *Trp53*^fl/fl^ (JAX #008462), were purchased from the Jackson Laboratory. *Rps19^fl/fl^*and *Rpl5^fl/fl^* were generated at The Jackson Laboratory as previously described(19).

All mice were housed under alternate dark/night cycle of 12h at 20 ± 2°C and 50 ± 10% humidity.

*Prrx-Cre^+^* male mice were used for breeding to avoid the known potential for germline allele excision when *Prrx1*-*Cre* is passed from the dam(21). Mouse tails were used for genotyping. The following primers were used for genotyping:

*Rps19* FORWARD (GAACGTGGGCTTGAGGGAGC);

*Rps19* REVERSE (GTTTGGCCAGCTTGACTGTG);

*Rpl5* WT FORWARD (AACCACCATGTAGTTGATGCC);

*Rpl5* WT REVERSE (GCTGACCGTCAATGCTTTCC);

*Rpl5* FLOX FORWARD (CCTAAACCACCATGTAGTTGATGATAACTTCGTATAATGTAT);

*Rpl5* FLOX REVERSE (CAGTGAGAAAAGTCAGGTTATTAACTTGTTGCTATCA);

*Prrx1-Cre* FORWARD (GCGGTCTGGCAGTAAAAACTATC);

*Prrx1-Cre* REVERSE (GTGAAACAGCATTGCTGTCACTT).

DNA extraction and isolation from tissues were performed according to manufacturer’s instructions (Invitrogen).

The study was reviewed and approved by the Feinstein Institutes IACUC, protocols #2022-019 and #25-0625.

### μCT analysis

Femurs were dissected of soft tissues and fixed in 70% ethanol at 4°C from 10-week-old male mice. Cortical morphometry of the femoral mid-diaphysis, and trabecular morphometry within the metaphyseal region of distal femurs, were quantified using X-ray micro-computed tomography (μCT50, Scanco Medical AG, Bassersdorf, Switzerland). Femurs were imaged in 70% ETOH at 70 kVp (200 μA), employing 2000 cone beam projections per revolution at an integration time of 500 msec. Three-dimensional images were reconstructed at ∼10 μm resolution using standard convolution back projection algorithms with Shepp and Logan filtering and rendered at a discrete voxel density of 1,000,000 voxels/mm^3^ (isometric 10 μm voxels). Bone mineral was calibrated to a discrete-step hydroxyapatite phantom and segmented from marrow and soft tissue in conjunction with a constrained Gaussian filter to reduce noise, applying mineral density thresholds of 700 and 500 mg HA/cm^3^ for cortical and trabecular bone, respectively. Anatomical boundaries of the diaphyseal cortex and the internal ROI within the trabecular compartment were identified using automated routines. Cortical morphometry was averaged within a mid-diaphyseal span of ∼0.7 mm (5% of femur length) to obtain measures of total area (Tt.Ar), cortical area (Ct.Ar), marrow area (Ma.Ar), polar Moment of Inertia (MOI), and the minimum and maximum dimensions of the mid diaphysis. Volumetric regions for trabecular bone analysis were selected within the endosteal borders to include the secondary spongiosa of femoral metaphyses located ∼0.7 mm (∼7% of length) from the growth plate and extending ∼1 mm proximally, scaling for bone length. Trabecular morphometric parameters were measured without imposing a presumed structural model to obtain direct measures of trabecular volume fraction (BV/TV), thickness (Tb.Th) and number (Tb.N, not shown).

### Whole mount skeletal staining

Whole skeletal preparations for cartilaginous and mineralized skeletal elements performed as described(36). Skeletal structures of newborn mice (P1) or 3-week old mice (P21) were dissected to remove the skin and fat and stained with Alizarin Red (Sigma-Aldrich A5533) for visualization of the mineralized elements and Alcian Blue (Sigma-Aldrich, B8438) to see the cartilaginous structures.

### Complete blood counts

Blood was collected from mice at P21 in EDTA-treated microcapillary tubes to prevent clot formation. Samples were analyzed using a Siemens ADVIA120 hematology system to obtain complete blood counts. No hemolysis affected the measurements.

### Bone immunofluoresence microscopy and histologic staining

Femurs and humeri bones were dissected and fixed in 4% paraformaldehyde (PFA) at 4°C overnight. After washing in water, samples were then decalcified for 10 days using 14% (w/v) EDTA (pH 7.4), embedded in paraffin, and sectioned onto slides at 5μm. Masson’s Trichrome stainings were performed according to manufacturer’s instructions. The sections were then counterstained with 0.1% Fast Green. Bright field imaging was performed on an EVOS M5000 Imaging System. Data are representative of at least three independent experiments.

### Mesenchymal stem cell (MSC) isolation, cell culture and differentiation

Mesenchymal stem cells were isolated from limbs of 3-4 weeks old mice of both sexes, according to (37). Bone chips were plated in 12-well and 6-well plates in MEM-A medium (GIBCO, 12561-056) supplemented with 10% FBS (Cytiva, SH30071.03), 1% penicillin-streptomycin (Thermo Scientific, 15140122) and cultured for 5-7 days at P0 before being passaged with Tryspin - EDTA 0.05% (Thermo Scientific, 25300054). Cells were passaged every 4-5 days until reaching P8. The culture media was replenished every 2 days.

### Flow cytometry analysis of MSCs and HSPCs

To assess MSC population, single cells were isolated from forelimbs and hindlimbs as described previously(25). Briefly, forelimbs and hindlimbs were isolated and cleaned from skin, skeletal muscle and connective tissues. Clean bones were then transferred to a mortar, digested (3,000 U/mL type II collagenase powder, 100 U/mL DNase, 1Mg/mL poloxamer 188 (P-188), 1 mg/mL BSA, 20 mM HEPES buffer, and 1 mM CaCl_2_ in Medium 199 with Hanks’ salts) and crushed. Minced bones were then incubated in digestion buffer for 30 min at 37 °C and filtered through a 75-p.m filter using at least 5 mL of cold cell-suspension buffer (2% (vol/vol) FBS, 1 mg/mL poloxamer (P-188), and 100 U/mL penicillin/streptomycin in calcium- and magnesium-free PBS). After centrifugation (5min, 500g), the cell pellet was resuspended in the same buffer and 10^6^ cells were stained with a cocktail of anti-CD45, Ter119, CD31, CD51 and Sca-1 conjugated antibodies. CD45^-^Ter119^-^CD31^-^CD51^+^Sca-1^+^ cells were gated and immunophenotypically defined as MSC.

To assess HSPC populations, 10^6^ cells were stained with lineage markers (CD4, CD8, Ter119, Ly6G/C, CD45R) and ckit. Lin^-^ckit^+^ cells were immunophenotypically defined as HSPCs.

### Protein synthesis measurements of bone-derived MSCs

To measure global protein synthesis, 10^6^ bone-derived cells were treated with 10μM O-Propargyl Puromycin (OPP) in DMEM (Gibco) medium for 10 min before harvesting. Cells were washed once in cold PBS and then stained with a cocktail of antibodies measuring MSCs (described above) for 30 min at RT. Cells were washed once (PBS, 2% FBS) and fixed with 1% PFA in PBS for 15 min at 4 °C. Cells were washed once (PBS, 2% FBS) and incubated in permeabilization buffer (PBS, 3% FBS, 0.1% saponin) for 5 min at RT. Click chemistry mix was prepared following the instructions of Click-iT^TM^ Edu Flow Cytometry Assay Kit (Invitrogen, C10632). Cells were incubated in Click chemistry mix at RT for 30 min and then washed twice with permeabilization buffer and twice with FACS buffer (PBS 0.5% BSA) before flow cytometry analysis.

### Protein synthesis measurements of cultured MSCs

MSC were isolated from the long bones as previously described(37) with the following modifications. MSCs at passage 1-2 isolated from 3-week-old mice were trypsinised, counted and resuspended at a concentration of 10^7^/ml on ice-cold PBS 0.1% BSA. After assessing MSC purity by flow cytometry, cells were washed 1x with 3ml ice – cold PBS 0.1% BSA and then fixed with 4% PFA in PBS for 15’ at RT. Cells were washed with 3ml ice-cold PBS 0.1% BSA and resuspended in 100μl ClickIT kit (Thermo Fisher Scientific C10269) permeabilization buffer and incubated for 15’ at RT before adding 400μl of the Plus reaction buffer of ClickIT kit, containing Alexa Fluor 555 Picolyl Azide dye (Thermo Fisher C10642), according to manufacturer’s instructions. Mix was incubated for 30’ at RT. Following the incubation, cells were washed 1x with 3ml permeabilization buffer, centrifuged and resuspended in 200μl of PBS 0.1% BSA and stained for 7-AAD (BD –Pharmingen 559925) for 10’ at RT protected from light before being analyzed in BD LSRFortessa Cell Analyzer. For fluorophore compensation OneComp eBeads from Invitrogen (01-1111-42) were used by mixing one drop of beads with 1μl of each antibody used in our panel while having unstained beads as control. The mix was incubated for 10’ at RT protected from light. 1ml of PBS 0.1% BSA and the mix was centrifuged and resuspended in 300μl PBS 0.1% BSA before analysis. For flow data analysis, Flowjo 10.5.3 software was utilized.

### Analysis of single-cell RNA sequencing

For scRNA-seq experiments, freshly isolated single cells from both forelimbs and hindlimbs at E17.5 were used. Library construction was performed with Chromium Next GEM Single Cell 3’ Library, Gel Bead & Multiplex Kit, and Chip Kit (10X Genomics) according to the manufacturer’s instructions. 3’ single-cell RNA-seq libraries generated with the 10X Genomics platform were processed using 10X Genomics Cell Ranger Suite (v7.0.0) with include-introns option, which aligned the data to mm10 reference genome, de-multiplexed and generated UMI count matrix files. Downstream analyses were performed in R (v4.5.0) using Seurat (v5.3.1). Seurat objects were generated from Cell Ranger-filtered matrices using a minimum of 200 detected features per cell and three cells per gene. The percentage of mitochondrial transcripts was calculated for each cell. Individual samples were normalized using SCTransform (v0.4.2) with the glmGamPoi method (v1.20.0), while regressing mitochondrial transcript percentage. Samples were integrated using the SCT reciprocal PCA (RPCA) workflow with 3,000 integration features and the first 30 principal components. UMAP dimensionality reduction and nearest-neighbor graph construction were performed using the first 20 principal components, followed by unsupervised clustering at a resolution of 1.0. A mitochondrial-high/low-quality cluster identified after clustering was excluded from downstream analyses. Cell populations were manually annotated based on the expression of established lineage-specific marker genes. Differential gene expression between mutant and WT cells was assessed separately for forelimbs and hindlimbs within each cell population using the Seurat FindMarkers function and Wilcoxon rank-sum test. Genes expressed in at least 10% of cells in either group were tested without an initial log-fold-change threshold. Genes with an adjusted P value <0.05 and |average log2 fold-change| >0.25 were considered differentially expressed. Gene set enrichment analysis was performed using fgsea (v1.34.2) and MSigDB gene sets obtained using msigdbr (v25.1.1). Genes were ranked according to their average log2 fold-change, and enrichment analyses were performed using Hallmark (H), curated (C2), Gene Ontology (C5), and immunologic signature (C7) gene sets. Trajectory and pseudotime analyses were performed using Slingshot (v2.16.0) on MSC/stromal and chondrocyte populations, including osteo/chondro progenitors, resting, proliferating, prehypertrophic, and hypertrophic chondrocytes. Slingshot was used to infer differentiation trajectories and assign pseudotime values along the mesenchymal-to-chondrocyte differentiation continuum.

### Statistics

Statistical analysis was performed using GraphPad Prism (v11). Unless otherwise specified, unpaired, two-tailed Student’s t test (single test) or one-way ANOVA (multiple comparisons) with a Tukey’s Post Hoc test were used to evaluate statistical significance (defined as *P<0.05*). All results are presented as mean value ± standard deviation. All measurements were taken from distinct samples, with a minimum of 3 biological replicates. No statistical method was used to predetermine sample size. These were based on our previous experience and published literature on similar models to ensure the robustness of the observed phenotypes.

### Study Approval

All human studies were conducted in accordance with the world medical association declaration of Helsinki. The fetal tissues were collected with informed and signed consent of the parents and under the supervision of the Institutional Review Board from the Hospital of St. Etienne, France and the ethical committee for biomedical research (CEERB-RD), Robert Debré hospital, Paris N°2012/005. The parents authorized the mutation screening analysis and the publication of the graphies of their fetus by signed informed consent.

### Data availability

All sequencing datasets have been deposited to NCBI Gene Expression Omnibus under accession code GSE353345. Any additional information with regards to the datasets is available from Dr. Lionel Blanc upon request. Source data are provided with this paper.

### Authorship

A.B., J.H. Y.T., L.L.P., L.D.C., J.M.L., C.A-B. and L.B. designed the research; A.B., J.H., Y.T., J.P., T.K., B.M.D., M.K., B.D., H.P., E.H., J.B., G.O., R.M., D.J.A., performed the experiments; M.E., R.T. and G.C-P. contributed to the analyses of the skeletal defects in the still born patient; R.R. contributed to the scRNAseq analyses, A.B., J.H., Y.T., J.P., T.K., B.M.D., H.P., J.B., G.O., R.M., D.A.G., P.M., B.J.B., A.N., N.M., I.R.., L.I.Z., S.D., L.L.P., L.D.C., J.M.L., C.A-B. and L.B. analyzed and interpreted the data; A.B., C.A-B. and L.B. cowrote the manuscript; and all authors read and commented on the final manuscript.

## Acknowledgments

The authors thank Dr. Raymond F. Robledo for technical assistance.

## Figure Legends

**Supplementary Figure 1.**
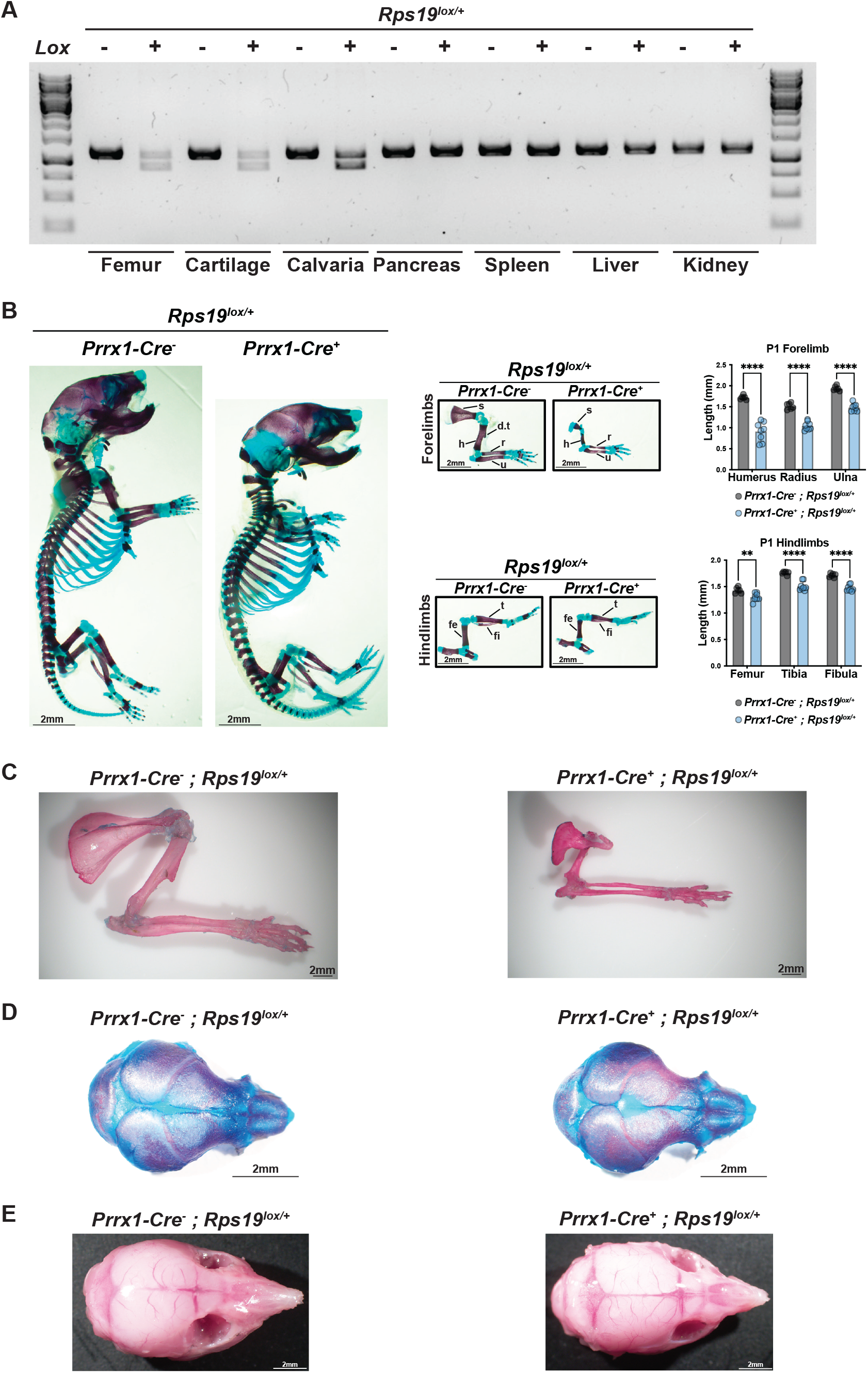
*Prrx1-Cre* promoter is specific to mesenchymal stem cells and leads to bone development defect. **A.** Genomic PCR for specific recombination of floxed Rps19 allele in tissues of osteogenic lineage compared to other tissues in *Prrx1-Cre^-^; Rps19^lox/+^* and *Prrx1-Cre^+^*; *Rps19^lox/+^* mice. **B.** Alcian Blue staining of *Prrx1-Cre^-^; Rps19^lox/+^* and *Prrx1-Cre^+^*; *Rps19^lox/+^* mice at P1. Red arrows denote the deltoid tuberosity. Quantification of length of forelimb and hindlimb bones. N=8 ± SD. Scapula (s), humeri (h), deltoid tuberosity (d.t), ulna (u), femur (fe), tibia (t), fibula (fi). *P*-values were calculated using multiple *t*-tests and adjusted for multiple comparisons using the Holm-Šídàk method. *\*\*P < 0.01; ****P < 0.0001.* **C.** Alizarin red staining of 10 weeks old hindlimb. **D.** Calvaria Alizarin S red and Alcian Blue staining of neonatal (P1) mice. **E.** Cranium pictures of three weeks old mice.

**Supplementary Figure 2.**
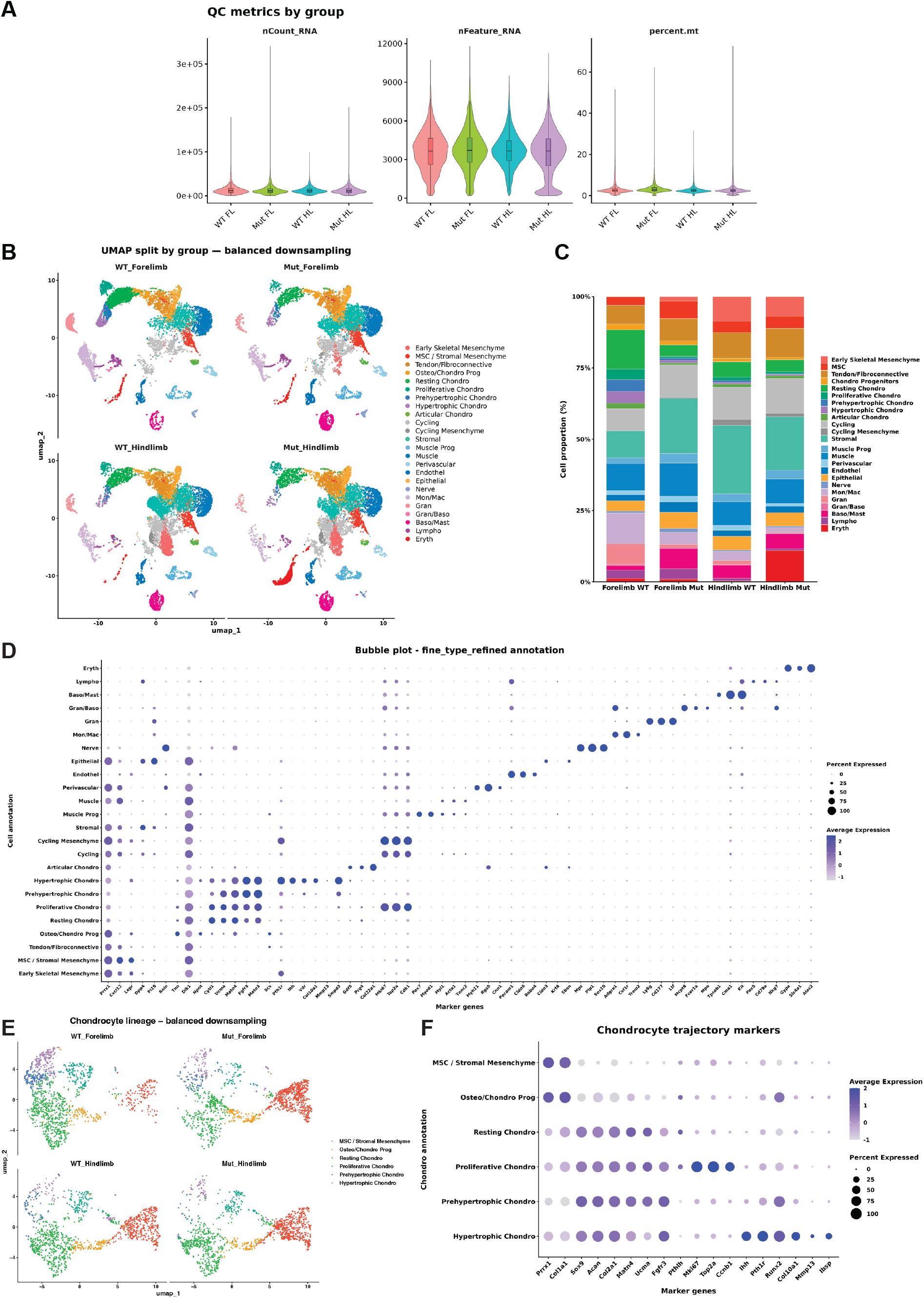
Single-cell transcriptomic characterization and annotation of limb cell populations. **A.** Quality-control metrics for the scRNA-seq datasets from WT and mutant forelimbs (FL) and hindlimbs (HL), showing the distribution of RNA counts (nCount_RNA), detected genes (nFeature_RNA), and mitochondrial transcript percentage (percent.mt) across groups. **B.** UMAP visualization of the integrated scRNA-seq dataset after balanced downsampling, split by genotype and limb. **C.** Relative abundance of the annotated cell populations in WT and mutant forelimbs and hindlimbs. Stacked bars represent the proportion of each cell type within each group. **D.** Dot plot showing expression of canonical marker genes used to define the major cell populations identified in the integrated dataset. Dot size represents the percentage of cells expressing each gene, and color indicates scaled average expression. **E.** UMAP visualization of the chondrocyte lineage after balanced downsampling, split by genotype and limb. **F.** Dot plot showing expression of representative markers associated with successive stages of chondrocyte differentiation across the six chondrocyte-lineage populations. Dot size represents the percentage of cells expressing each gene, and color indicates scaled average expression.

**Supplementary Figure 3.**
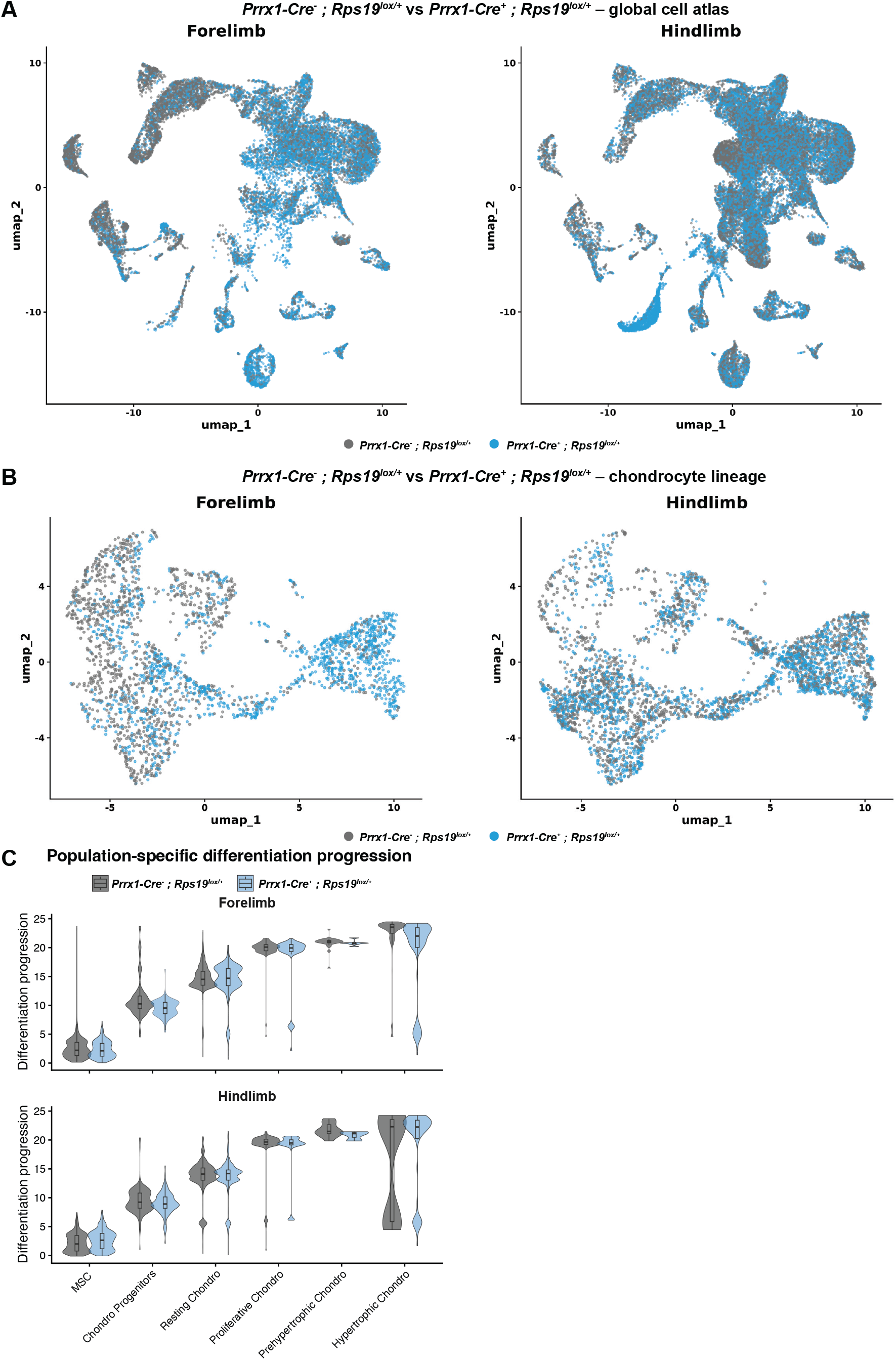
Rps19 haploinsufficiency alters the distribution of cells along the chondrocyte differentiation trajectory. **A.** UMAP overlays of WT and mutant cells across the global limb cell atlas in forelimbs (left) and hindlimbs (right). WT and mutant cells are shown in gray and blue, respectively. **B.** UMAP overlays of WT and mutant cells restricted to the chondrocyte lineage in forelimbs (left) and hindlimbs (right). WT and mutant cells are shown in gray and blue, respectively. **C.** Distribution of the differentiation progression score across successive chondrocyte-lineage populations in forelimbs (top) and hindlimbs (bottom). Violin plots show WT and mutant cells within each population.

**Supplementary Figure 4.**
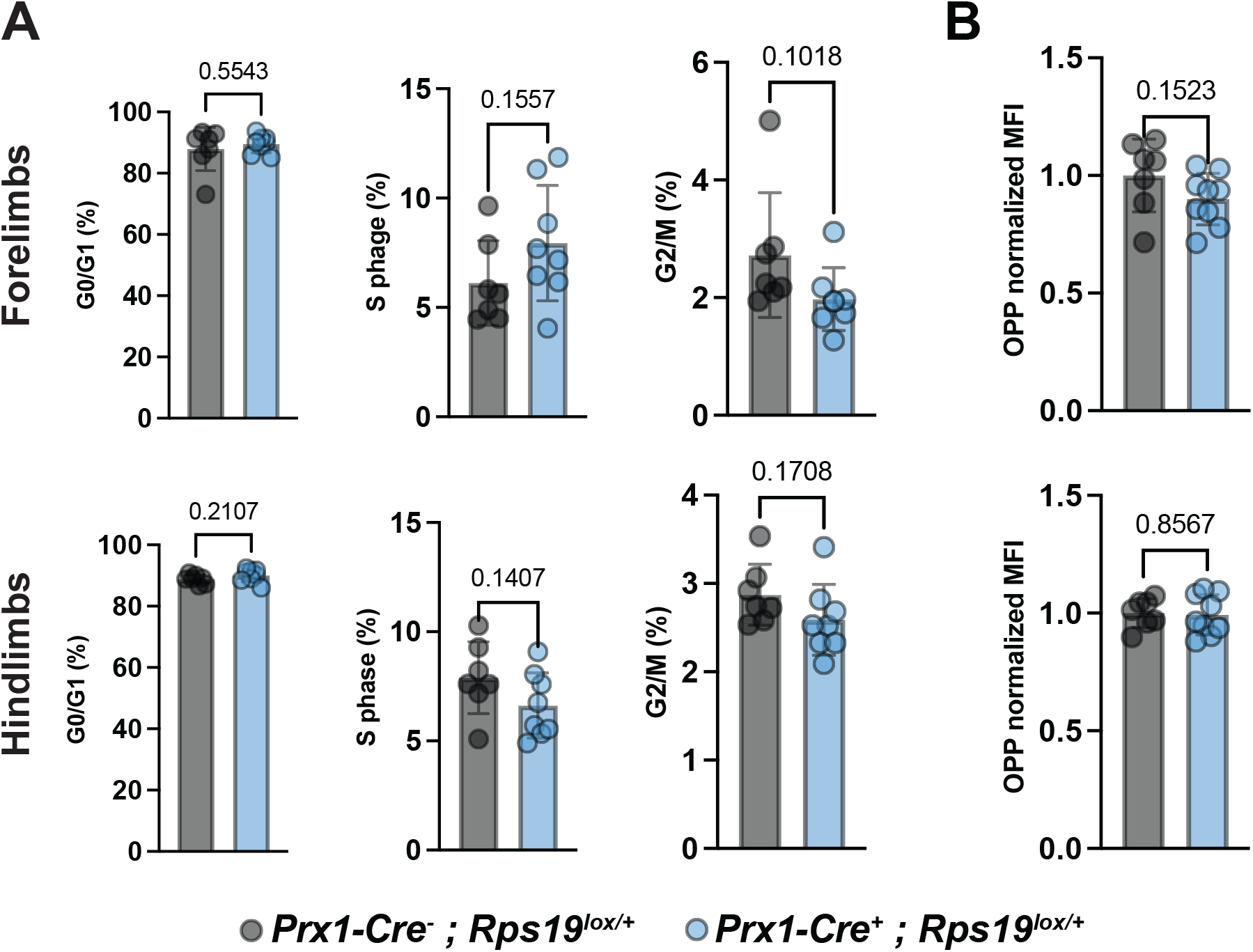
*In vivo* Cell cycle distribution and protein synthesis are not altered in the mutant MSCs. **A.** Cell-cycle distribution of MSCs from control (*Prrx1^-^*) and mutant (*Prrx1^+^*) *Rps19^lox/+^* E17.5 forelimbs (top) and hindlimbs (bottom), showing the percentage of cells in G0/G1, S, and G2/M phases. **B.** Protein synthesis in forelimb (top) and hindlimb (bottom) MSCs assessed by O-propargyl-puromycin (OPP) incorporation. OPP mean fluorescence intensity (MFI) was normalized to the control group.

**Supplementary Figure 5.**
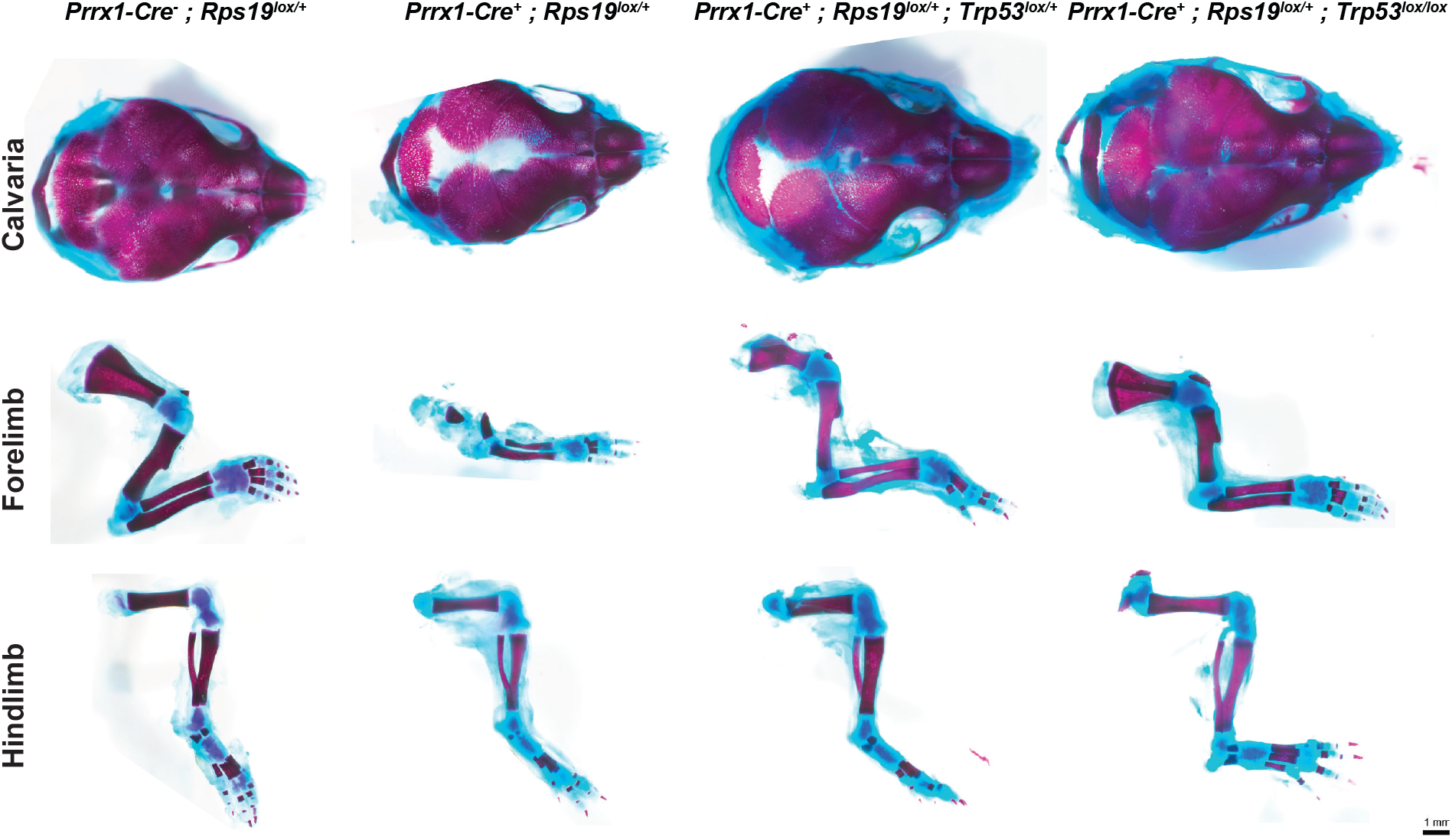
**Skeletal phenotype and p53-dependent rescue of Rps19 haploinsufficiency**

## REFERENCES

1. Genuth NR, and Barna M. The Discovery of Ribosome Heterogeneity and Its Implications for Gene Regulation and Organismal Life. Mol Cell. 2018;71(3):364–74.

2. Harnett D, Ambrozkiewicz MC, Zinnall U, Rusanova A, Borisova E, Drescher AN, et al. A critical period of translational control during brain development at codon resolution. Nat Struct Mol Biol. 2022;29(12):1277–90.

3. Mercer M, Jang S, Ni C, and Buszczak M. The Dynamic Regulation of mRNA Translation and Ribosome Biogenesis During Germ Cell Development and Reproductive Aging. Front Cell Dev Biol. 2021;9:710186.

4. Tiu GC, Kerr CH, Forester CM, Krishnarao PS, Rosenblatt HD, Raj N, et al. A p53-dependent translational program directs tissue-selective phenotypes in a model of ribosomopathies. Dev Cell. 2021;56(14):2089–102 e11.

5. Xue S, and Barna M. Specialized ribosomes: a new frontier in gene regulation and organismal biology. Nat Rev Mol Cell Biol. 2012;13(6):355–69.

6. Shi Z, and Barna M. Translating the genome in time and space: specialized ribosomes, RNA regulons, and RNA-binding proteins. Annu Rev Cell Dev Biol. 2015;31:31–54.

7. Gay DM, Lund AH, and Jansson MD. Translational control through ribosome heterogeneity and functional specialization. Trends Biochem Sci. 2021.

8. Narla A, and Ebert BL. Ribosomopathies: human disorders of ribosome dysfunction. Blood. 2010;115(16):3196–205.

9. Da Costa L, Leblanc T, and Mohandas N. Diamond-Blackfan anemia. Blood. 2020;136(11):1262–73.

10. Ulirsch JC, Verboon JM, Kazerounian S, Guo MH, Yuan D, Ludwig LS, et al. The Genetic Landscape of Diamond-Blackfan Anemia. Am J Hum Genet. 2018;103(6):930–47.

11. James A, Wang Y, Raje H, Rosby R, and DiMario P. Nucleolar stress with and without p53. Nucleus. 2014;5(5):402–26.

12. Dutt S, Narla A, Lin K, Mullally A, Abayasekara N, Megerdichian C, et al. Haploinsufficiency for ribosomal protein genes causes selective activation of p53 in human erythroid progenitor cells. Blood. 2011;117(9):2567–76.

13. Vlachos A, Blanc L, and Lipton JM. Diamond Blackfan anemia: a model for the translational approach to understanding human disease. Expert Rev Hematol. 2014;7(3):359–72.

14. Gazda HT, Sheen MR, Vlachos A, Choesmel V, O’Donohue MF, Schneider H, et al. Ribosomal protein L5 and L11 mutations are associated with cleft palate and abnormal thumbs in Diamond-Blackfan anemia patients. Am J Hum Genet. 2008;83(6):769–80.

15. Vlachos A, Rosenberg PS, Atsidaftos E, Kang J, Onel K, Sharaf RN, et al. Increased risk of colon cancer and osteogenic sarcoma in Diamond-Blackfan anemia. Blood. 2018;132(20):2205–8.

16. Calvi LM, and Link DC. The hematopoietic stem cell niche in homeostasis and disease. Blood. 2015;126(22):2443–51.

17. Yu VW, and Scadden DT. Heterogeneity of the bone marrow niche. Curr Opin Hematol. 2016;23(4):331–8.

18. Sieff C. In: Adam MP, Ardinger HH, Pagon RA, Wallace SE, Bean LJH, Mirzaa G, et al. eds. GeneReviews((R)). Seattle (WA); 1993.

19. Tang Y, Ling T, Mehmood R, Bertrand A, Papoin J, Khan M, et al. RPS19 and RPL5 haploinsufficient models reveal divergent ribosomal subunit controls of fetal hematopoiesis. Nat Commun. 2026;17(1).

20. Da Costa L, Chanoz-Poulard G, Simansour M, French M, Bouvier R, Prieur F, et al. First de novo mutation in RPS19 gene as the cause of hydrops fetalis in Diamond-Blackfan anemia. Am J Hematol. 2013;88(2):160.

21. Logan M, Martin JF, Nagy A, Lobe C, Olson EN, and Tabin CJ. Expression of Cre Recombinase in the developing mouse limb bud driven by a Prxl enhancer. Genesis. 2002;33(2):77–80.

22. Glatt V, Canalis E, Stadmeyer L, and Bouxsein ML. Age-related changes in trabecular architecture differ in female and male C57BL/6J mice. J Bone Miner Res. 2007;22(8):1197–207.

23. Rowe DW, Adams DJ, Hong SH, Zhang C, Shin DG, Renata Rydzik C, et al. Screening Gene Knockout Mice for Variation in Bone Mass: Analysis by muCT and Histomorphometry. Curr Osteoporos Rep. 2018;16(2):77–94.

24. Durland JL, Sferlazzo M, Logan M, and Burke AC. Visualizing the lateral somitic frontier in the Prx1Cre transgenic mouse. J Anat. 2008;212(5):590–602.

25. Pisterzi P, Chen L, van Dijk C, Wevers MJW, Bindels EJM, and Raaijmakers M. Resource: A Cellular Developmental Taxonomy of the Bone Marrow Mesenchymal Stem Cell Population in Mice. Hemasphere. 2023;7(2):e823.

26. Lu L, Harutyunyan K, Jin W, Wu J, Yang T, Chen Y, et al. RECǪL4 Regulates p53 Function In Vivo During Skeletogenesis. J Bone Miner Res. 2015;30(6):1077–89.

27. Ǫuarato ER, Salama NA, and Calvi LM. Interplay Between Skeletal and Hematopoietic Cells in the Bone Marrow Microenvironment in Homeostasis and Aging. Curr Osteoporos Rep. 2024;22(4):416–32.

28. Signer RA, Magee JA, Salic A, and Morrison SJ. Haematopoietic stem cells require a highly regulated protein synthesis rate. Nature. 2014;509(7498):49–54.

29. Da Costa L, Mohandas N, David NL, Platon J, Marie I, O’Donohue MF, et al. Diamond-Blackfan anemia, the archetype of ribosomopathy: How distinct is it from the other constitutional ribosomopathies? Blood Cells Mol Dis. 2024;106:102838.

30. Moniz H, Gastou M, Leblanc T, Hurtaud C, Cretien A, Lecluse Y, et al. Primary hematopoietic cells from DBA patients with mutations in RPL11 and RPS19 genes exhibit distinct erythroid phenotype in vitro. Cell Death Dis. 2012;3(7):e356.

31. Komori T, Yagi H, Nomura S, Yamaguchi A, Sasaki K, Deguchi K, et al. Targeted disruption of Cbfa1 results in a complete lack of bone formation owing to maturational arrest of osteoblasts. Cell. 1997;89(5):755–64.

32. Lee B, Thirunavukkarasu K, Zhou L, Pastore L, Baldini A, Hecht J, et al. Missense mutations abolishing DNA binding of the osteoblast-specific transcription factor OSF2/CBFA1 in cleidocranial dysplasia. Nat Genet. 1997;16(3):307–10.

33. Lengner CJ, Steinman HA, Gagnon J, Smith TW, Henderson JE, Kream BE, et al. Osteoblast differentiation and skeletal development are regulated by Mdm2-p53 signaling. J Cell Biol. 2006;172(6):909–21.

34. Vlachos A, Osorio DS, Atsidaftos E, Kang J, Lababidi ML, Seiden HS, et al. Increased Prevalence of Congenital Heart Disease in Children With Diamond Blackfan Anemia Suggests Unrecognized Diamond Blackfan Anemia as a Cause of Congenital Heart Disease in the General Population: A Report of the Diamond Blackfan Anemia Registry. Circ Genom Precis Med. 2018;11(5):e002044.

35. Iskander D, Wang G, Heuston EF, Christodoulidou C, Psaila B, Ponnusamy K, et al. Single-cell profiling of human bone marrow progenitors reveals mechanisms of failing erythropoiesis in Diamond-Blackfan anemia. Sci Transl Med. 2021;13(610):eabf0113.

36. Rigueur D, and Lyons KM. Whole-mount skeletal staining. Methods Mol Biol. 2014;1130:113–21.

37. Zhu H, Guo ZK, Jiang XX, Li H, Wang XY, Yao HY, et al. A protocol for isolation and culture of mesenchymal stem cells from mouse compact bone. Nat Protoc. 2010;5(3):550–60.

